# DNA Methylation Signatures in Breast Cancer: A Systematic Review and Meta-Analysis

**DOI:** 10.1101/2022.10.15.512358

**Authors:** Antonio Manuel Trasierras-Fresco, Z Andreu, Beatriz Dolader Rabinad, Marta R. Hidalgo, Ricardo A. Hernández-Ramírez, Irene Soler-Sáez, Borja Gómez-Cabañes, Atocha Romero, Jose A Lopez-Guerrero, María de la Iglesia-Vaya, Francisco García-García

## Abstract

Epigenetic changes are involved in the onset and progression of cancer, and the detection of DNA methylation signatures may foster the improvement of diagnosis and prognosis. While the emergence of innovative technologies has fostered numerous studies in breast cancer, many lack statistical power due to the small sample sizes generally involved. In this study, we present a novel meta-analysis that identifies a common pattern of DNA methylation in all breast cancer subtypes. We obtained DNA methylation signatures at the gene and biological function level, identifying those significant groups of genes and functional pathways affected. To achieve this, we conducted a thorough systematic review following PRISMA statement guidelines for the selection of studies on DNA methylation in breast cancer. In total, we gathered four studies (GSE52865, GSE141338, GSE59901 and GSE101443) that were split into 13 comparisons comprising a set of 144 individuals. We discovered that most breast cancer subtypes share a significant deregulation in the immune system and alterations to the cell cycle. This integrative approach combines all available information from public data repositories and possesses greater statistical power than any individual study. Further evaluations of the identified differential biological processes and pathways may support the identification of novel biomarkers and therapeutic targets.

**Simple summary:** The identification of DNA methylation patterns in breast cancer represents a potentially valuable approach in defining more accurate diagnoses and treatment options. In this study, we applied a novel methodology that integrates the DNA methylation profiles of all studies available in public repositories via systematic review and meta-analysis. The results provide evidence of a common DNA methylation signature in distinct breast cancer subtypes, which reflects a significant deregulation of the immune system and alterations to the cell cycle. Overall, these results may support the selection of disease/treatment biomarkers and the identification of therapeutic targets.

## 1. Introduction

Despite ongoing advances in research into disease mechanisms and treatment approaches, breast cancer remains a major health problem that affects millions of women worldwide. Breast cancer represents the leading cause of cancer-related deaths in women aged <45 years due to the development of metastases, with incidence and mortality rates both expected to increase significantly in the coming years [78]. The highly heterogeneous and potentially aggressive nature of breast cancer may derive from the aberrant accumulation of epigenetic changes, which can determine the effectiveness of various treatments [1].

Transcriptomic studies of breast cancer have revealed the presence of different molecular subtypes with vastly different prognoses [79]. Efforts that aim to improve the understanding of breast cancer have focused on: i) early detection mechanisms, such as the identification of hormone receptor status [2], ii) the development of monoclonal antibody therapies, which can treat approximately 30% of breast cancers [3], and iii) the development of technologies to evaluate gene expression profiled of affected patients. A more in-depth understanding of the mechanisms controlling breast cancer development and spread [1] may foster the development of better diagnostic tools to allow early detection and improved and personalized treatment approaches that may reduce mortality.

DNA methylation represents a major epigenetic mechanisms in mammals that generally acts to repress gene expression by inducing a repressive or “closed” chromatin environment and inhibiting the binding of transcription factors to gene regulatory regions. Abnormal DNA methylation patterns contribute to several diseases including cancer [4] - abnormal hypermethylation of tumor suppressor genes and hypomethylation of oncogenes can permit the uncontrolled proliferation of cancer cells, leading to tumorigenesis.

High-throughput -omic technologies such as microarrays and next generation sequencing have enabled major advances in cancer research. The huge amounts of cancer genomic data generated is housed in open access biological repositories, such as the Gene Expression Omnibus (GEO) ^1^, Array Express ^2^, and Sequence Read Archive (SRA) ^3^. Such databases represent an important source of biological information that can be reanalyzed with different approaches and interests in mind.

While the cost of high-throughput technologies has decreased significantly in recent times, sample size remains a limiting factor in cancer research, leading to the problem of class-imbalanced datasets that hampers correct statistical analysis of the data [5]. Meta-analysis, which allows a measure of the combined effect of interest and offers greater precision than individual studies, may overcome this problem. Meta-analysis and related techniques have been applied in numerous fields such as social sciences [6], medicine [7] and genomics [8], where methods frequently focus on the gene [9–15] or variant level [16,17].

In this study we present a novel targeted approach at two complementary levels: gene and biological function, which integrates the different genes or regions that present significant biological activity, thus providing a better understanding of the mechanisms of the disease in a systems biology framework. The main objective of the study is to detect and characterize epigenetic biomarkers and those mechanisms altered in breast cancer that may aid diagnosis and prognosis. For this purpose, we undertook a functional meta-analysis [18] to identify DNA methylation signatures associated with various subtypes of breast cancer (for the first time, to the best of our knowledge). We hope that this approach will support the identification of new biomarkers for breast cancer subtypes that may aid the development of diagnostic/predictive tools for breast cancer management and novel treatment approaches.

## 2. Results

We have organized the results into five sections.

i) the selection of studies from the systematic review
ii) and iii) results of the bioinformatic analysis for each selected study, taking in the exploratory analysis, differential methylation and functional characterization.
vi) and v) a description of the main results, which includes the methylation profiles at the gene and function level identified via meta-analysis.

We have made the detailed results of each section available in the metafun-BC web tool (https://bioinfo.cipf.es/metafun-BC), which allows the user to review the results described in the manuscript.

### 2.1. Systematic Review and Selection of Studies

Following the inclusion criteria, we identified and carefully reviewed a total of 77 studies to avoid duplicates and ensure a focus on breast cancer, which finally provided 39 full-text articles. We retrieved raw DNA methylation data from five studies (Figure 1 - GSE78751, GSE52865, GSE141338, GSE101443, and GSE59901), which represents a total of 14 different comparisons between case and control groups including a total of 185 patients (150 cases and 35 controls) (Figure 2). Adjacent to tumor normal breast tissue was considered (paired samples) in all the studies unless GSE59901 where it is not indicated.

**Figure 1.**
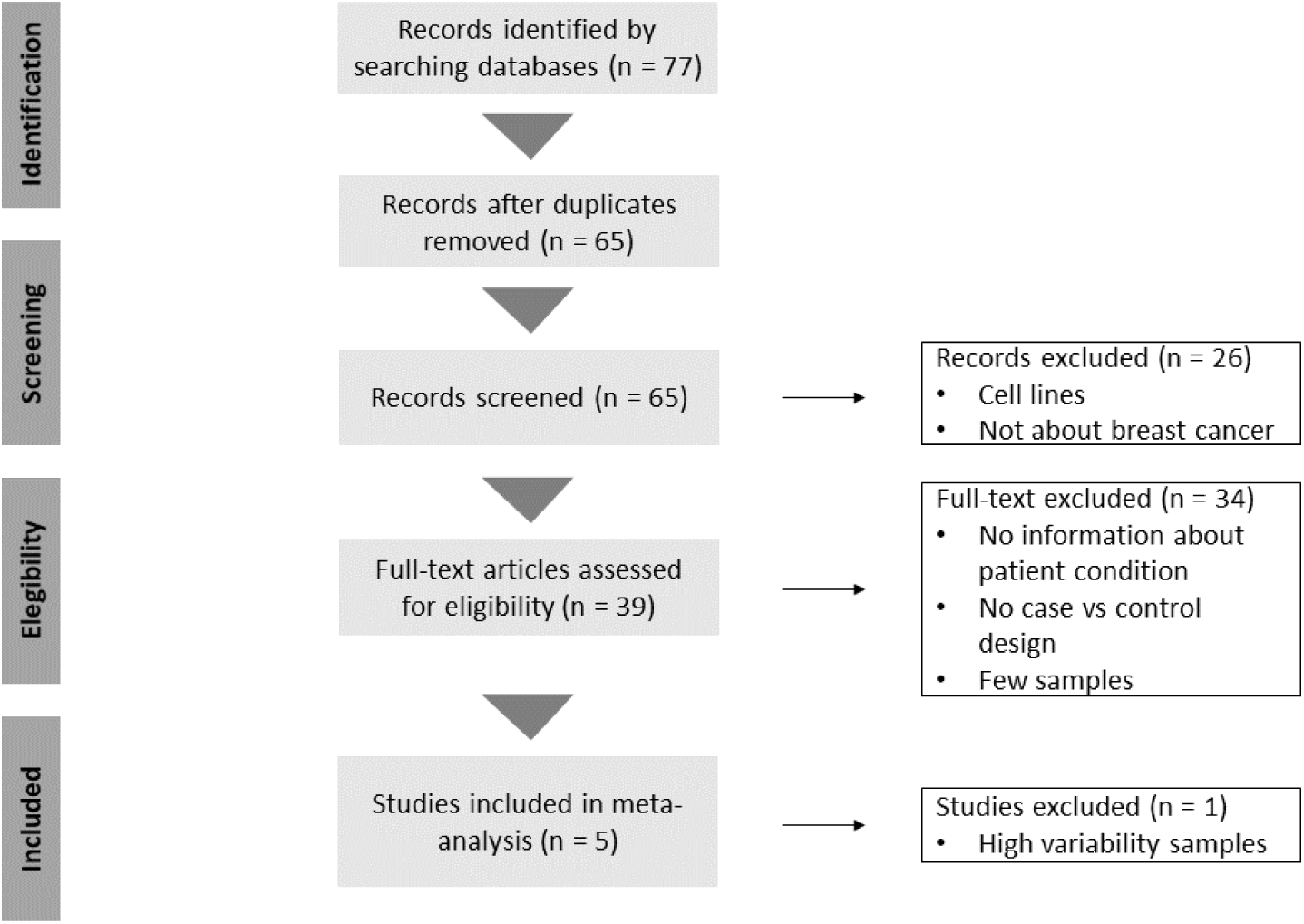
Flow diagram presenting our systematic review of the literature and selection of studies for this meta-analysis, according to the PRISMA statement guidelines.

**Figure 2.**
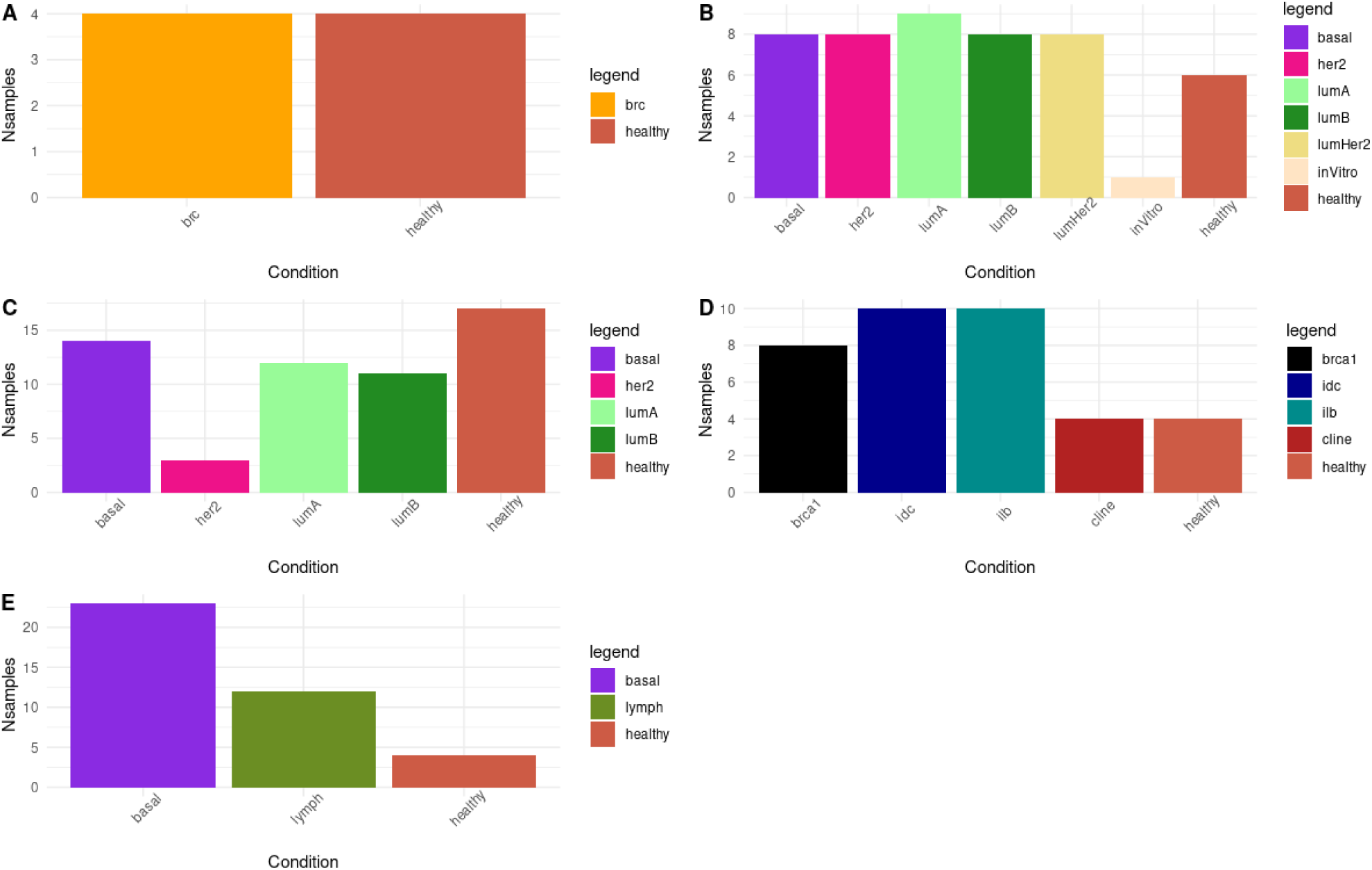
Studies selected for meta-analysis. **A, B, C, D**, and **E** report the distribution of samples for studies GSE101413 [19], GSE141338 [21], GSE52865 [22], GSE59901, and GSE78751 [23] respectively. Abbreviations: brc - no subtype specified, healthy – healthy breast tissue, basal - triple negative/basal-like subtype, her2 - her2-positive subtype, lumA -- luminal A subtype, lumB - luminal B subtype, lumHer2 - luminal-Her2 undifferentiated, inVitro - in vitro samples, brca1 - BRCA1-mutated breast cancer, idc - invasive ductal carcinoma, ilb - invasive lobular carcinoma, and lymph - lymph node metastasis.

The GSE101443 study (A) comprises four tumor samples and four healthy samples, which we included in the next step of the meta-analysis. For the GSE141338 study (B), only healthy and tumor samples were considered for the meta-analysis (41 cancer and 6 healthy samples). In the case of the GSE59901 (D) and GSE78751 (E) studies, we discarded information regarding cell lines and lymph node metastasis (4 and 15 samples, respectively) and we considered only the breast tissues (28 tumor and 4 healthy samples for the GSE59901 study and 23 tumor and 4 healthy samples in the case of the GSE78751 study). Finally, we considered all samples from the GSE52865 (C) study (40 tumor and 17 healthy samples). Figure 2 presents a detailed configuration of the different datasets.

### 2.2. Data Exploration, Quality Control, and Normalization

We explored all datasets by principal component analysis (PCA) and clustering analysis (see the “Analysis exploratory” section in the metafun-BC web tool). We failed to detect any abnormal patterns except for the GSE78751 study, which presented with high variability. To avoid this negative influence on the outcome of the subsequent meta-analysis [20], we chose to discard this study.

### 2.3. Individual Epigenomic Analysis

We applied the same workflow analysis to all individual studies and collected the analysis of differentially methylated genes in the form of UpSet plots. Figure 3 shows (for each study) the number of significant genes and specific and common differentially methylation status according to the methylation score (MS): higher MS (Figure 3a) and lower MS (Figure 3b). We found highly variable results for the differential methylation analysis for each of the 13 comparisons evaluated. In 10 comparisons, the number of hypomethylated genes in the breast cancer samples varied between 0 and 423, whereas the hypermethylated genes ranged between 1 and 39 (Figure 3). Interestingly, we found a common pattern of methylated genes in breast cancer samples between different studies. For example, Figure 3b shows how the luminal-A and luminal-B subtypes from the GSE52865 study share a total of 69 hypomethylated genes and 3 hypermethylated compared to control samples (see Figure 3a).

**Figure 3.**
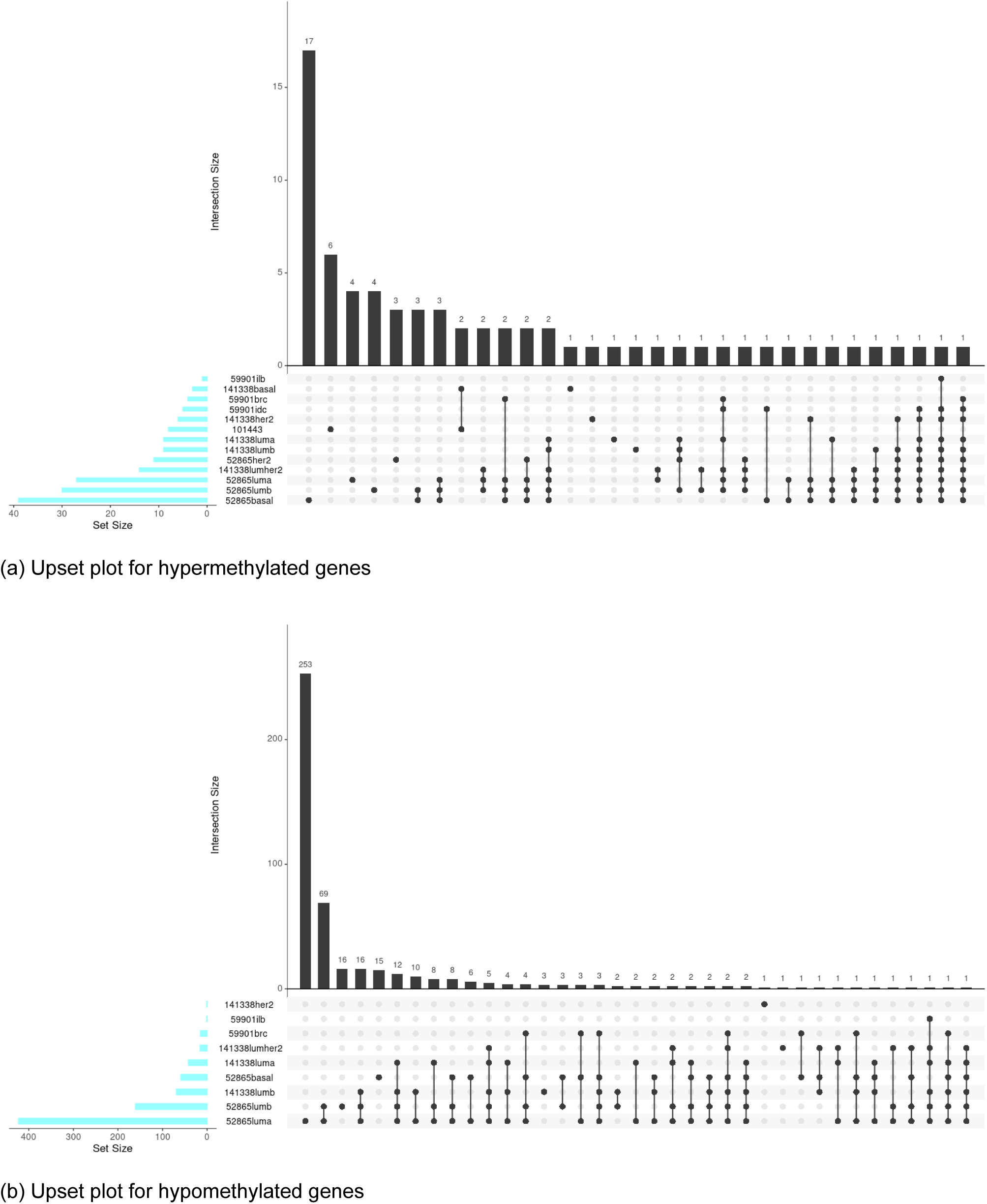
UpSet plots showing the number of common elements among the significantly hypermethylated genes (a) and hypomethylated genes (b) in the differential methylation analysis. Horizontal bars indicate the number of significant elements in each study. The vertical bars indicate the elements in common between the sets indicated with dots under each bar. The unique points represent the number of unique elements in each group.

Of note, UpSet plots demonstrate a lack of significant genes across all studies. This data highlights the need for integrated strategies, such as meta-analyses, to increase the statistical power of any findings.

For each of the comparisons described, we functionally profiled the list of all genes ranked by their differential MS. Table 1 shows the GSEA results for the Gene Ontology terms and KEGG pathways – the intersection of significant results failed to highlight common functions across the different studies.

**Table 1.** Number of significant GO terms and KEGG pathways from GSEA. Sig Up and Sig Down refer to the GO terms or KEGG pathways that are significantly hypermethylated and hypomethylated, respectively, in breast cancer samples.

|  | GO BP |  | GO MF |  | GO CC |  | KEGG |  |
| --- | --- | --- | --- | --- | --- | --- | --- | --- |
|  | Sig Up | Sig Down | Sig Up | Sig Down | Sig Up | Sig Down | Sig Up | Sig Down |
| 52865-basal | 25 | 28 | 2 | 2 | 1 | 0 | 0 | 1 |
| 52865-her2 | 14 | 58 | 0 | 1 | 1 | 0 | 0 | 1 |
| 52865-luma | 31 | 14 | 3 | 8 | 19 | 5 | 1 | 3 |
| 52865-lumb | 34 | 16 | 3 | 2 | 0 | 0 | 0 | 1 |
| 59901-brc | 44 | 25 | 4 | 15 | 11 | 1 | 1 | 1 |
| 59901-idc | 45 | 11 | 7 | 5 | 7 | 0 | 0 | 1 |
| 59901-ilc | 24 | 9 | 7 | 6 | 0 | 0 | 0 | 1 |
| 101443-bc | 12 | 6 | 7 | 3 | 5 | 5 | 1 | 1 |
| 141338-basal | 24 | 12 | 15 | 2 | 28 | 13 | 2 | 1 |
| 141338-her2 | 26 | 5 | 1 | 3 | 1 | 0 | 2 | 2 |
| 141338-luma | 3 | 28 | 2 | 18 | 7 | 18 | 3 | 2 |
| 141338-lumb | 8 | 17 | 4 | 10 | 0 | 8 | 0 | 1 |
| 141338-lumher2 | 7 | 7 | 2 | 1 | 0 | 0 | 0 | 1 |

We integrated those results in the final functional meta-analysis in the hope of retrieving common functions affected in the different breast cancer studies.

### 2.4. Methylated Gene Meta-Analysis

The meta-analysis strategy provided 19 protein-coding and micro-(mi)RNA genes with a common methylation profile in all comparisons - 4 genes displayed hypermethylation (*INMT-MINDY4*, *LTB4R2*, *VSTM2B* and *ZNF471*), 13 genes (*FAM25C, GABRA5, HSFY1P1, HULC, MAGEA5, NKAPP1, OR1B1, OR1Q1, RPS16P5, TGIF2LY, CCL16, CCL3, NREP-AS1*) displayed hypomethylation, and 2 microRNAs (*hsa-miR-122* and *hsa-miR-384*) displayed hypomethylation compared to controls (Table 2).

**Table 2.** Final set of significantly affected protein-coding and miRNA genes obtained by meta-analysis. P-value (adjusted by Benjamini-Hochberg) and methylation pattern are shown for each protein-coding or miRNA gene. + indicates hypermethylation and – indicates hypomethylation compared to controls.

| Gene / miRNA | P-value | P-adj | Methylation |
| --- | --- | --- | --- |
| CCL16 | 0.00298 | 0.04164 | - |
| ZNF471 | 0.00279 | 0.02282 | + |
| INMT-MINDY4 | 0.00324 | 0.01768 | + |
| MIR384 | 0.00108 | 0.01518 | - |
| NREP-AS1 | 0.00094 | 0.01321 | - |
| GABRA5 | 0.00146 | 0.01023 | - |
| CCL3 | 0.00019 | 0.00259 | - |
| FAM25C | 0.00008 | 0.00114 | - |
| NKAPP1 | 0.00007 | 0.00102 | - |
| TGIF2LY | <0.00001 | 0.00002 | - |
| LTB4R2 | <0.00001 | <0.00001 | + |
| VSTM2B | <0.00001 | <0.00001 | + |
| HSFY1P1 | <0.00001 | <0.00001 | - |
| HULC | <0.00001 | <0.00001 | - |
| MAGEA5 | <0.00001 | <0.00001 | - |
| MIR122 | <0.00001 | <0.00001 | - |
| OR1B1 | <0.00001 | <0.00001 | - |
| OR1Q1 | <0.00001 | <0.00001 | - |
| RPS16P5 | <0.00001 | <0.00001 | - |

### 2.5. Functional DNA Methylation Signature Meta-Analysis

The functional meta-analysis integration of distinct functions retrieved from the individual studies represents the last step of the workflow. We considered a random effects mode to account for the possible heterogeneity in the effect measurements in the different studies. Table 3 summarizes the significant GO terms and KEGG pathways that we uncovered.

**Table 3.** Significant Results of the Meta-analysis. GO:BP, GO:MF, and GO:CC refer to GO terms related to biological processes, molecular functions, and cellular components, respectively. KEGG refers to KEGG pathways. Functions with an adjusted p-value less than or equal to 0.05 were considered significant.

| DB | Sig Up | Sig Down | No Sig |
| --- | --- | --- | --- |
| GO:BP | 1187 | 819 | 6824 |
| GO:MF | 185 | 134 | 1456 |
| GO:CC | 115 | 66 | 933 |
| KEGG | 28 | 19 | 175 |

We conducted a total of 11,941 meta-analyses for the GO terms and KEGG pathways considered in the previous individual analyses (11,719 and 222 meta-analyses, respectively). The results of the functional meta-analysis show the magnitude of the combined effect of all the individual studies through the logarithm of the odds ratio (LOR). We identified a large number of significant functions; detailed results are available in the Functional Meta-analysis section of metafun-BC web tool.

We next focused on the GO biological processes and KEGG pathways obtained that displayed significantly different DNA methylation profiles and exhibited a high magnitude of change (LOR >= -0.5 or LOR >= 0.5). Thus, we obtained a total number of significant GO biological processes obtained of 138 (85 hypermethylated and 53 hypomethylated), while no significant KEGG pathway passed the filtering criteria. Table 4 and Figure 4 show a selection of these biological processes with higher magnitude of change.

**Figure 4.**
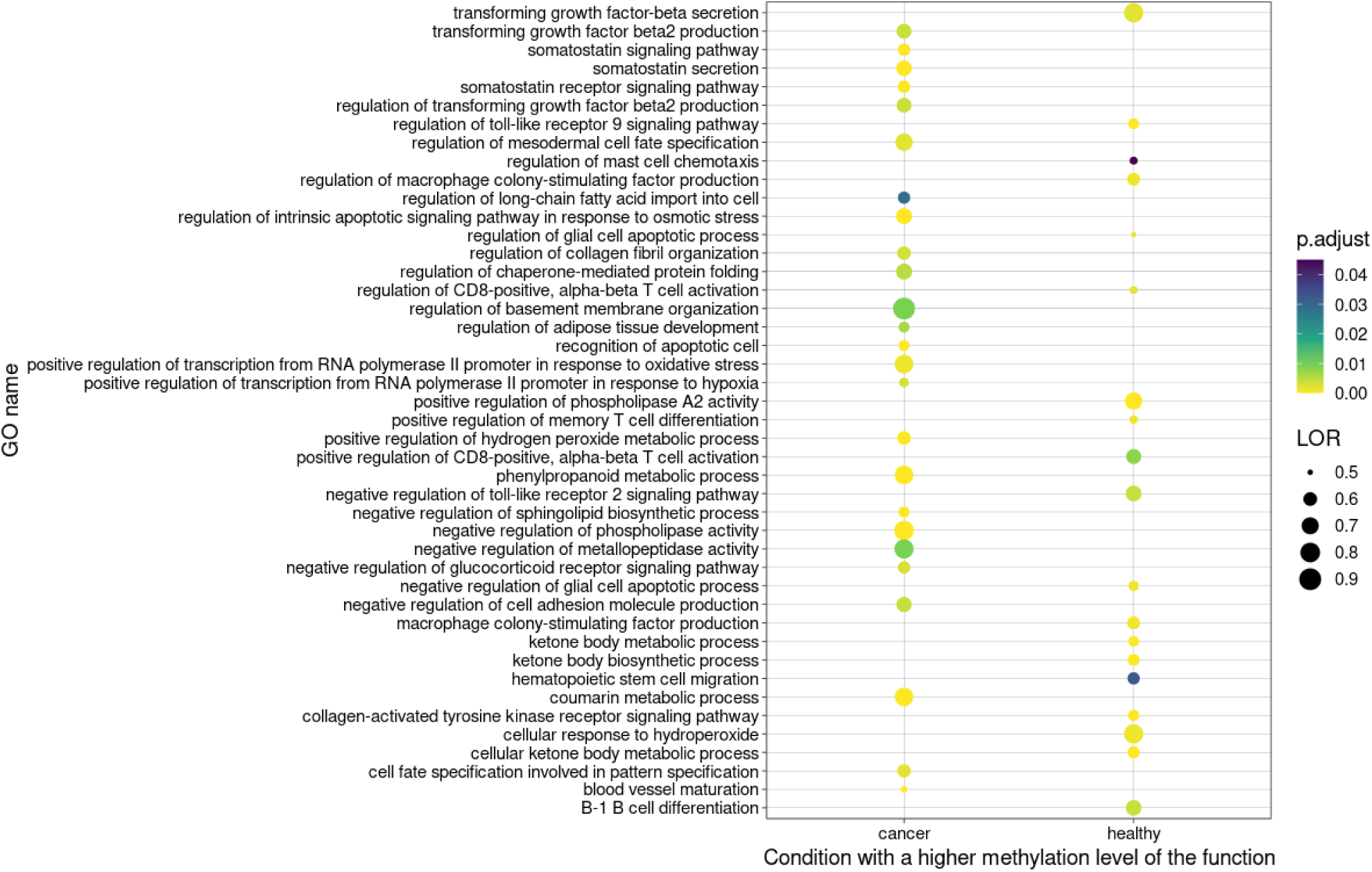
Dotplot of selected significant functions obtained by functional meta-analysis. The thickness of the circle refers to the number of genes related to a specific pathway. The GeneRatio refers to the number of genes related to a pathway relative to the total number of genes in the functional enrichment analysis. The adjusted p-values are represented by a color gradient.

**Table 4.**
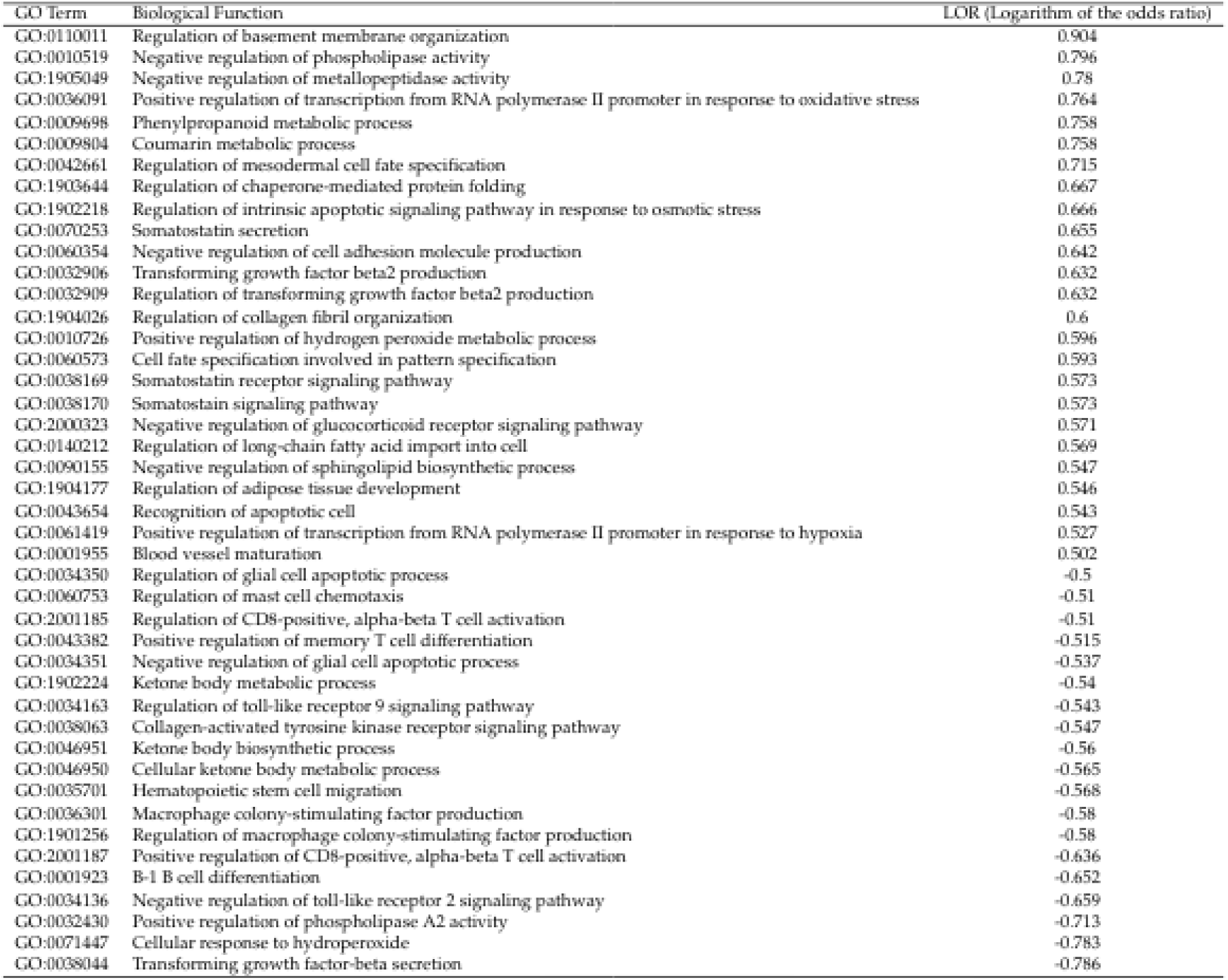
List of functions related to Figure 4. A positive LOR indicates hypermethylation in breast cancer samples and a negative LOR indicates hypomethylation in breast cancer samples when compared to healthy control samples.

## 3. Discussion

In this study, we generated a new characterization of breast cancer based on DNA methylation patterns. This novel strategy based on the meta-analysis of reviewed and selected studies (whose data are available in public repositories) has provided new DNA methylation signatures at the gene and biological function level. These results supported the identification of common markers across breast cancer subtypes that may impact the identification of new diagnostic/predictive tools and the development of potential therapeutic approaches.

### 3.1. Gene-Level Meta-Analysis

Differentially DNA methylated regions detected in the breast cancer samples identified in this study suggest the silencing of the *INMT-MINDY4, LTB4R2, VSTM2B,* and *ZNF471* genes. Little is known about *INMT-MINDY4* and *VSTM2B* in cancer.

INMT-MINDY4 is a lysine48 deubiquitinase 4 that belongs to a lncRNA gene category described as regulatory molecules of gene expression. This gene has been related with FAM188B (family with sequence similarity 188, member B), an oncogenic protein highly expressed in most solid tumors as lung and colorectal cancer that negatively correlates with the overall survival of lung cancer patients. *INMT-MINDY4-FAM188B* locus represents a rare but naturally occurring read-through transcription between *INMT-MINDY4* and *FAM188B* on chromosome 7. The read-through transcript is unlikely to produce a protein. Interestingly, siFAM188B treatment induced the upregulation and activation of *TP53*, and consequently increased p53-regulated pro-apoptotic protein oncogenic role [25, 26]. The hypermethylation of the *VSTM2B* gene has been previously reported in human papillomavirus-related oropharyngeal squamous cell carcinoma [27]. *LTB4R2*, a leukotriene receptor also known as *BLT2*, is associated with malignant cell transformation in esophageal squamous cell carcinoma and pancreatic cancers [28, 29], invasion, metastasis, and survival in TNBC, and paclitaxel resistance in breast cancer cell lines [30]. The silencing of *VSTM2B* and *LTB4R2* might suggest a better prognosis for breast cancer patients. *ZNF471* codes for a zinc finger protein, whose members have previously shown tumor suppressor activity and high rates of DNA methylation in primary head and neck squamous cell carcinoma according to our analysis [27, 31]. In breast cancer, ZNF471 exerts a tumor-suppressive function by blocking AKT and Wnt/β-catenin signaling pathways; furthermore, downregulated expression through epigenetic regulation has been associated with worse survival [32].

Hypomethylated genes include *CCL16* and *CCL3*, cytokines whose overexpression can contribute to breast cancer progression and metastases [33, 34]. We also encountered the hypomethylation of the *FAM25* and *GABRA5* genes, suggesting their overexpression. While there exist no know link between FAM25 and cancer, studies have suggested a role in the brain and in drug resistance (Allen Brain Atlas Adult Human Brain Tissue Gene Expression Profiles). Meanwhile elevated expression of GABRA5 has been related to pediatric brain cancer medulloblastoma. Although further studies must be performed, our results open the possibility to consider *FAM25* and *GABRA5* as biomarkers for brain metastases, a common complication in TNBC, and of poor prognostic. Expression of the *HULC* gene, also hypomethylated in our study, can prompt the epithelial-mesenchymal transition to promote tumorigenesis and metastases in hepatocellular carcinoma. The hypomethylated *MAGEA5* gene has been described as tumor suppressor in breast cancer (comparably to other MAGEA genes) whose expression associates with improved relapse-free survival [36].

Our study also showed the hypomethylation of regions associated with *miR-122* and *miR-384*, suggesting their expression. The upregulated expression of *miR-122* inhibits cell proliferation and suppresses tumorigenesis in vivo by targeting IGF1R in breast cancer [37]. Additionally, miR-122 also differentially controls the response to radiotherapy through a dual function as a tumor suppressor and an oncomiR, dependent on cell phenotype [38]. The overexpression of *miR-384* has been demonstrated to inhibit the proliferation and migration of breast cancer cells in vitro and in vivo. Therefore, the presence of hypomethylation at the *miR-122* and *miR-384* genes (or their overexpression) could represent prognostic biomarkers for breast cancer patients and help to predict response to therapy.

The dysregulation of pseudogene expression in human cancer has been linked to tumor initiation and progression [39], and we identified hypomethylated regions related to the *HSFY1P1, NKAAP1,* and *RPS16P5* pseudogenes in this study. HSFY1P1 is responsible of cat eye syndrome [40], and while the biological functions of HSFY genes are poorly understood, accumulating evidence links the overexpression of heat shock factors in a variety of human cancer to growth, survival, and metastasis [41].

Our methylation analysis also highlighted the hypomethylation of **NREP-AS1**, an antisense RNA associated with the NREP gene, which has been reported as a predictive biomarker for prostate cancer progression [42]. We also found hypomethylation of *OR1B1* and *OR1Q1*, which encode olfactory receptors, in breast cancer samples. Mutations in these genes have been associated with neoplasms [43].

The proposed integrative meta-analysis has identified differentially methylated regions (DMRs) related to expression genes with potential utility as prognostic indicators, therapeutic targets and guiding treatment decisions.

### 3.2. Functional-Level Meta-Analysis

Our study identified an elevated number of deregulated functions common to all breast cancer studies, which relate to well-known cancer hallmarks, such as apoptosis, immune response, and metastases [44] Table 4 lists the top functions (GO BP) collected in Figure 4, ordered according to their level of differential methylation in breast cancer samples.

Apoptosis may be deliberated by via extrinseca mediated by receptors as tumor necrosis factor (TNF) and intrinsic pathways in response to DNA damage with the participation of mitochondria and mitochondrial proteins [45], in addition to T-cell mediated cytotoxicity and perforin-granzyme system. But frequently cancer cells show resistance to apoptosis programs [46]. Related to apoptosis, we found altered the biological processes GO:0043654 (the recognition of apoptotic cell) and GO:1902218 (the regulation of intrinsic apoptotic signaling pathway in response to osmotic stress), both driving resistance to apoptosis in breast cancer cells [45]. In our study, genomic regions involved in the recognition of apoptotic cell and intrinsic pathway processes present an elevated level of methylation in breast cancer patients that means a loss or lower level of the expression of the genes involved in apoptotic mechanisms and suggests in these breast cancer patients the apoptosis evasion by tumor cells. Additionally, tumor cells of these patients could be able to evade traditional therapies such as chemotherapy, radio, and immunotherapy since resistance to apoptosis may confer resistance to conventional therapies and the immune system. The Programmed cell death is a known cancer hallmark, critical to the maintenance of genomic homeostasis. Thus, and according to the literature [47] truncation of the apoptotic signaling pathway by several factors such as DNA damage or osmotic stress play a critical role in the development of several cancers. In fact, as we noted from our results, there is not only a clear avoidance of the cell death program, but also a dysregulation of functions related to stress response. Among other functions obtained, the regulation of chaperone-mediated protein folding (GO:1903644) and functions related to hydrogen peroxide species stand out in this regard. It is common for cancer cells to be subjected to a wide variety of types of stress, including oxidative stress, DNA damage, hypoxia, and nutrient deficiencies, among others.

Many of these stress signals culminate in programmed cell death in favor of the rest of the healthy tissue, but mutations in cancer can lead proteins such as chaperones to protect cancer cells from programmed death. Chaperones are a family of proteins involved in the folding of proteins so that they can perform their function. There is great evidence that they are found in high concentration in distinct types of cancer. This is because, for cancer cells to avoid programmed cell death, they require chaperones to stabilize poorly folded proteins due to these elevated levels of stress, which could otherwise lead to cell death [48]. On the other hand, our results suggest a higher level of expression of functions related to the cellular response to oxidative stress as we can note by the GO Term 0071447 (Cellular response to hydroperoxide). This function is found with lower methylation levels in cancer samples which means that it is highly expressed in these samples. Hydroperoxide is an important signaling molecule that is produced by several types of cancer. This molecule could increase genomic instability by inducing damage to DNA strands, which could facilitate the appearance of malignant processes such as proliferation, resistance to apoptosis and metastasis among others. However, other studies indicate that a relatively high concentration of this molecule may be able to induce selective apoptosis in cancer cells [49].

Overall, the response to stress and avoidance of cell apoptosis observed in the study could lead to an improved overall survival of breast cancer cells which is a recurrent characteristic denoted by many types of cancer.

It is also worth noting that several of our results also agree with a higher proliferative profile of breast cancer cells in contrast with healthy ones, which indeed is another major cancer hallmark. In this sense, we found a deregulation of the genes that carry out the biological process GO:0032430 which refers to the positive regulation of phospholipase A2 activity. These genes are found with a lower level of methylation in cancer patients, which results, a priori, in a higher level of expression of the genes which carry out this function. According to the literature [50], phospholipase A2 enzymes regulate the release of biologically active fatty acids and lysophospholipids from membrane phospholipid pools. In general, phospholipases are important mediators in intracellular and intercellular signals. The lipids generated by these proteins can act by promoting tumorigenesis [51], or modulating proliferation, migration, invasion, and angiogenesis. Although the protein phospholipase A2 has not yet been classified as an oncogene or tumor suppressor gene, there is compelling evidence that their key role in breast cancer could be more related with oncogenic functions.

Moreover, we can find other biological processes such as GO terms GO:0032906 and GO:0038044, both related to the transforming growth factor beta protein (TGF-B). Although these results may seem divergent (note that one is over methylated and the other under methylated, resulting in a higher and lower expression respectively), it is interesting to remark that the dual effects of TGF-B during tumor growth in breast cancer have been reported in the literature. In fact, many discordant results have been published about its prognosis in breast cancer. Among them, the consensus is that TGF-B influences cell homeostasis through proliferation, migration, and apoptosis. TGF-B has been reported to function as a tumor suppressor in the initial stages of the disease by inhibiting cell proliferation, however, it appears that in later stages it would have pro-oncogenic capabilities through stimulation of cell invasion and migration [52].

It is important to note the presence of other terms associated with a high proliferative state of breast cancer cells such as the somatostatin signaling pathway (GO terms GO:0038169 and GO:0038170). Somatostatin receptors initiate a signaling cascade that increases apoptosis and represses cell proliferation. According to our results, genes belonging to this pathway are less expressed in individuals with breast cancer (higher methylation status on cancer samples). This is an expected result since routes that lead to an apoptotic anti-proliferation signal are often silenced in several types of cancer. However, according to several studies [53, 54] somatostatin receptors are found in large numbers on breast cancer cells. This seems to point out that in our studies, some point on the somatostatin signaling pathway may be affected. According to the literature cited above, it is possible that somatostatin activates tumor suppressor genes such as PTEN and p53, so it appears that it may be this point on the somatostatin signaling pathway that is found to be under-expressed in our samples. Even so, treatment with somatostatin is usually as effective that research lines have been opened to fight cancer through its analogues. This is a signalling pathway that would be worth studying further to find out which exact point in the pathway might be affected. Last but not least for this functional block of functions let us discuss the term GO:0090155 which makes reference to the negative regulation of sphingolipid biosynthetic processes. Evidence that lipid mediators play pivotal roles in breast cancer biology has been increasing for the last few years. Since that, sphingolipids have emerged as important signaling mediators that regulate critical processes during the development of cancer. Silencing of genes related with this term could be related with a higher production of sphingolipids in breast cancer tissue with respect to normal breast tissue which indeed is a behavior reported in the literature [55]. In this way, high sphingolipids levels may be related to cancer proliferation and metastasis in breast cancer cells.

According to the results obtained in Table 4, let us now consider a group of functions related to vascularization and tissue invasion, which is another important hallmark in cancer intimately linked to tumor dissemination. Within this group of functions, we find the regulation of mesodermal cell fate specification (GO:0042661). This biological process is linked to a deregulation of the mesoderm, which could point to one of the main characteristics of several types of tumors: the epithelial to mesenchymal transition (EMT). Due to EMTs, cancer cells acquire a strong invasive and metastatic capacity. In addition, it has been shown that there is a certain capacity of modulation, being a reversible process. This great invasive capacity is given because mesenchymal cells are separated from each other by their respective cellular matrices, not having a basal lamina that separates them from the adjacent tissue and therefore having greater freedom of movement [56].

On the other hand, related to a higher capacity of invasion we found functions related to the structural organization of the cells. For example, let us consider the function regulation of basement membrane organization (GO:0110011). The basement membrane is an important part of the extracellular matrix which underlies epithelial and endothelial tissues. This structure serves as a natural barrier against cancer invasion, intravasation and extravasation, however, it is known that cancer cells can invade it to spread across other areas [57]. In fact, we obtained other functions related with the invasion and migration of cancer cells through the extracellular matrix. Among these functions, note the downregulation of the organization of the collagen fibers (term GO:1904026, caused by a high methylation level in cancer samples) and a high activity of metallopeptidases (GO:1905049) caused by a downregulation of genes related to the negative regulation of their activity. First, the downregulation of collagen fibers has been widely described during years on every phase of carcinogenesis and tumor progression. Changes in the organization and structure of collagen fibers contribute to the formation of a microenvironment that promotes cancer progression and invasion [58]. Related to the tumoral microenvironment, the increased activity of metallopeptidases could also favor cancer invasion and migration. Metallopeptidases are a family of proteins which have been recently proposed as markers of many cancers due to their ability to degrade extracellular matrix components and remodel tissues. Some studies have pointed out that the overexpression of these proteins can lead to a loss of epithelial phenotype and the adoption of mesenchymal one, increasing the migratory capabilities of cancer cells [59]. Overall, with our results we observe a high invasive and migratory status on breast cancer cells.

Metabolism disruption is another key factor for the survival of cancer cells. Through a readjustment of the metabolism, cancer cells obtain selective advantages during initiation and progression such as deregulated uptake of glucose and amino acids or opportunistic modes of nutrient acquisition [60]. There is a high evidence of the important role of lipids in cancer progression because they are required as a structural part of the cell membranes, to provide energy to the cell or just as secondary messengers [61]. This phenomenon can be seen in our results with GO BPs like the regulation of long-chain fatty acid import into cell (GO:0140212). The genes responsible for this function appear with higher methylation levels in cancer samples which means that their expression could be silenced in this condition. This kind of metabolism deregulation has been widely observed in several types of cancer because, as stated before, cancer cells need nutrients in order to keep growing and proliferation. In this way, the lipid metabolism is not only affected by a higher uptake of peripheral lipids into the cell but also by a deregulation in the development of the adipose tissue (GO:1904177) which ultimately led to an increased lipid metabolism in breast cancer cells. On the other hand, related to cancer metabolism we obtained the terms GO:0046951 and GO:0046950 which refer to ketone body metabolic process and ketone body biosynthetic process. According to the results, both functions show a lower methylation profile in cancer samples than in healthy ones which means that the expression of the genes belonging to the related functions is higher in cancer condition. Not only lipid metabolism but ketone metabolism also has been reported to be overexpressed in breast cancer cells. Ketones have a two-compartment metabolism in tumor cells [62]. First by upregulating key enzymes in the production of ketogenic fibroblasts in the stroma of adjacent breast cancer cells. In a second step, these ketone bodies are transferred from stromal fibroblast to cancer cells. Through this metabolic process breast cancer cells will obtain energy for promoting tumor growth and metastasis.

Finally, a growing evidence on the implication of the immune system with cancer has been reported in recent years. In this way, let us focus on a group of functions related to the immune system and immune modulation in breast cancer samples. Note that all functions related to the immune system at Table 4 are found with a lower level of methylation in breast cancer patients than in healthy individuals. Within this functional block we find the following GO terms: GO:0060753, GO:2001185, GO:0043382, GO:0034351,GO:0034163, GO:0035701, GO:0036301, GO:1901256, GO:2001187, GO:0001923 and GO:0034136. The Immune system is intimately involved in the progression of cancer through various functions such as its pro-inflammatory activity [63]. For example, the activation of T cells — GO:2001187 — is a process that presents a lower level of methylation in cancer patients, which leads to a higher level of expression of genes related to this function. T cells can have both pro-inflammatory and anti-inflammatory activities while their infiltration into cancerous tissues correlates with a better prognosis of the disease. However, it has been observed that several types of cancer are able to utilize the immunosuppressive properties of T cells and in turn, alter the anti-tumor effects they exert, such as their infiltration, survival, proliferation and cytotoxicity capabilities. In this way, cancer cells that escape the control of the immune system adopt a phenotype that is resistant to the immune system while taking advantage of its pro-inflammatory capabilities.

On the other hand, some cancers have a lower amount of leukocyte antigens on their surface due to several mutations. Thanks to this phenomenon, cancer cells can avoid the effect of leukocytes on them while their infiltration causes a higher rate of inflammation, which they take advantage of to develop [64]. It is also worth noting the terms GO:0036301 and GO:1901256 referring to a higher methylation level of genes related to the macrophage colony-stimulating factor production. Tumor associated macrophages can promote cancer progression in several ways, for example by secreting IL-6 which enhances cancer epithelial-to-mesenchymal transition through p-STAT3 signaling [65]. Additionally, macrophages can contribute to cancer progression by altering glucose metabolism, promoting angiogenesis and immune evasion within the tumor [66]. Finally we found a deregulation of genes related to toll-like receptors 2 and 9 signaling pathways (GO:0034136 and GO:0034163 respectively).

Toll-like receptors (TLRs) are pattern recognition receptors which can be found on the surface of immune cells. Their main function is the production of cytokines and chemokines to promote the inflammatory response. Although the roles of TLRs have not been widely studied in breast cancer yet, there is evidence that supports the crosstalk between them and breast cancer. In our results, we found that the negative regulation of the TLR 2 has higher methylation levels on normal breast tissues which could be related with a lower expression level of the genes responsible for carrying out that function. In contrast, in breast cancer cells we would find the opposite, a higher expression of the genes which repress this signaling pathway. Comparable results have been reported in the literature [67]. In this case, the expression levels of the TLR 2 were about 10-fold lower in the breast cancer cell line MDA-MB-231 compared to the less malignant MCF7 breast cancer cells. However, there seems to be controversy regarding the role of this type of receptor in breast cancer since the overactivation of TLR 2 signaling pathway promotes upregulation of interleukin-6, transforming growth factor-B, vascular endothelial growth factor and matrix metalloproteinase 9. On the other hand, TLR 9 has been positively correlated with tumor grade suggesting that this receptor is related to poor differentiated breast cancer tissue and involved in tumor progression and metastasis. In this sense, it is the only toll-like receptor (along with TLR 6) whose expression is associated with greater tumor aggressiveness. Overall, although promising, further studies are needed to fully elucidate the role and mechanism of action of TLRs in breast cancer. As we can see, our results agree with what is expected in the literature: the immune system is intimately linked to cancer and may even favor its development and invasiveness through inflammatory processes. Thus, the up regulation of the immune system functions in our study point to the creation of an ideal tumor microenvironment for the development of the disease and could guide breast cancer immunotherapy.

This dual characterization of methylation profiles in breast cancer has provided which elements could be relevant in a broad set of subtypes of this disease, identifying a gene signature as well as a function signature. This information provides guidance on the main molecular mechanisms that may be the target of new therapeutic targets.

## 4. Materials and Methods

### 4.1. Systematic Review and Selection of Studies

This review was conducted in October 2019, according to the Preferred Reporting Items for Systematic Reviews and Meta-Analyses (PRISMA) statement guidelines [68]. Breast cancer methylation datasets were selected from GEO repository [69], using the following keywords: *breast cancer*, *methylation,* and *Homo sapiens* in studies published in English. The following exclusion criteria were applied: (i) studies conducted in organisms other than humans; (ii) sample size less than 10 in each experimental group; (iii) experimental design different from case-control and (iv) methylation profiling platform other than Infinium HumanMethylation450 BeadChip from Illumina. The study platform was chosen for its high resolution - around 450,000 methylation sites - and its wide acceptance in the field [19].

### 4.2. Bioinformatics Analysis Strategy

The following workflow was applied to each of the selected studies: i) Data acquisition; ii) Exploratory analysis and quality control of the samples; iii) splitting of the studies into different *case vs. control* comparisons; iv) Analysis of differential methylation profiles between case and control groups; v) functional enrichment analysis for each comparison; and, vi) integration of the methylation profiles and functional results in the final meta-analysis —see Figure 5A—.

**Figure 5.**
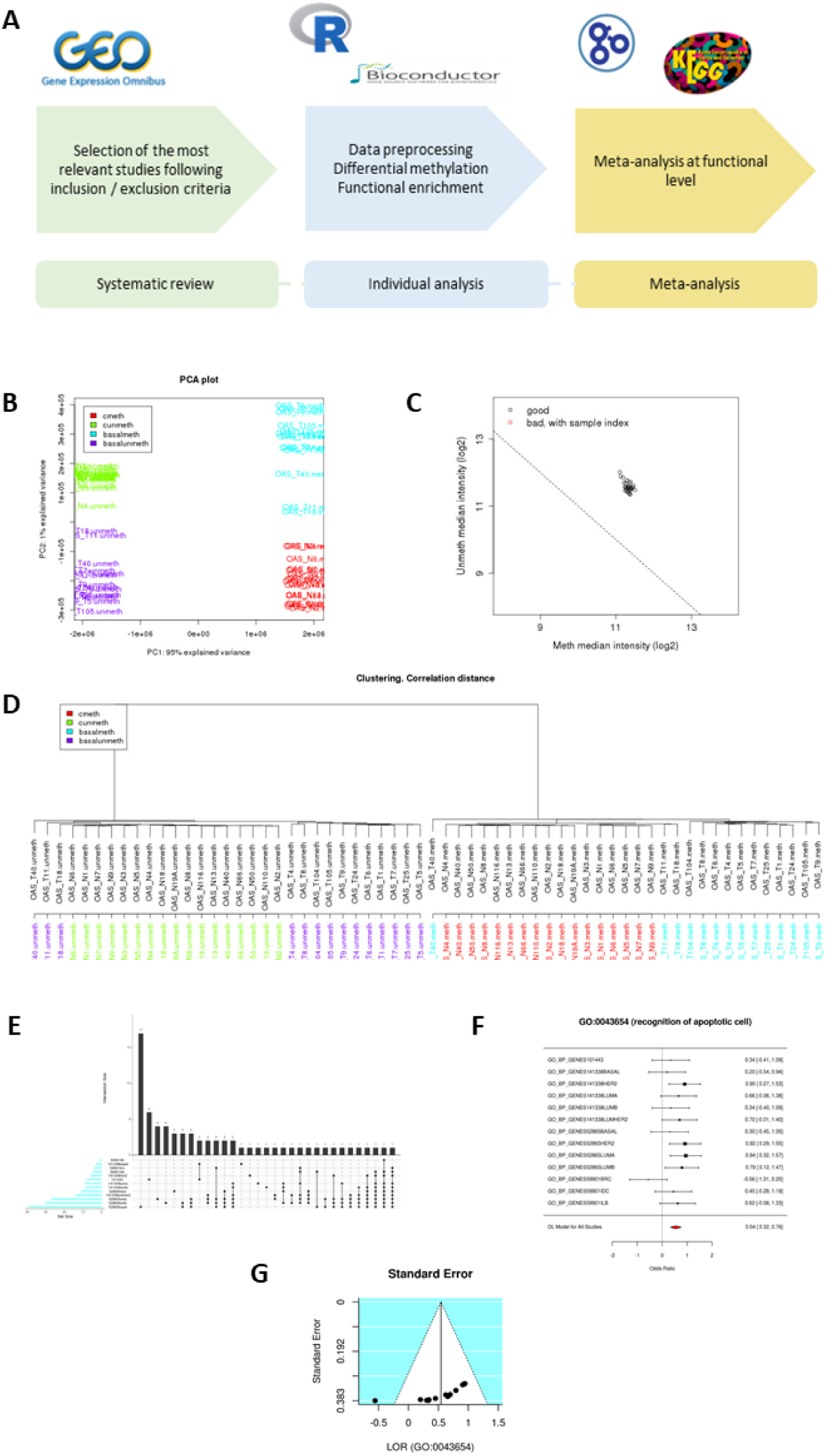
(A) Pipeline data analysis. (B, C and D) Exploratory data analysis. (E) UpSet plot showing the number of common elements among the significant genes. Only the 20 most abundant interactions are shown. Horizontal bars indicate the number of significant elements in each study. The vertical bars indicate the common elements in the sets, indicated with dots under each bar. The single points represent the number of unique elements in each group. (F) A forest plot of the GO:0042110 term, showing the LOR of each study and the global result. (G) Funnel plot of the GO:0042110 term; dots in the white area indicate the absence of bias and heterogeneity.

### 4.3. Data Exploration, Quality Control and Normalization

Exploration of raw data was performed through principal component analysis (PCA) and clustering analysis. Data quality control was conducted with the minfi R package [20]. Levels of the signals from methylated and non-methylated channels were checked, then a quantile normalization was performed on the data for subsequent comparisons —Figures 5B, 5C, 5D—.

### 4.4. Individual Epigenomic Analysis

Differential methylation analysis for each comparison was carried out with the minfi R package. Briefly, permutation tests were performed to obtain the methylation scores of all analyzed regions on the Illumina BeadChip 450k. A total of 13 comparisons derived from the individual studies were performed in the form *control vs cancer* (where a positive methylation score indicates a higher methylation value on cancer samples than in the control condition). Comparisons were performed between paired samples for all the studies except for the GSE59901 study where the information about the source of the control samples were missing. The comparisons are: Control vs TNBC (Triple Negative Breast Cancer) (GSE52865 and GSE141338), Control vs HER-2 subtype (GSE52865 and GSE141338), Control vs Lum-A subtype (Luminal A) (GSE52865 and GSE141338), Control vs Lum-B subtype (Luminal B) (GSE52865 and GSE141338), Control vs *BRCA1 mutated* (no subtype specified), Control vs Invasive Ductal Carcinoma (IDC), Control Vs Invasive Lobular Carcinoma (ILC) and Control vs BRCA (no molecular subtype specified).

Subsequently, methylation scores were annotated to gene level through the bumphunter R Package [70]. In cases where more than one differentially methylated region (DMR) was reported for a gene, the one with the highest absolute value was used. Then, a functional enrichment analysis was performed from the differential methylation results using Gene Set Analysis (GSA) [71] in which the genes were ordered according to their p-values and the sign of the contrast statistic. The GSA was then performed following the logistic regression model implemented in the mdgsa R package [72], as well as its corresponding functional annotation. The p-values were corrected for each function by false discovery rate (FDR) [73]. The databases used for functional enrichment were the Gene Ontology (GO) [74] and the Kyoto PATHWAY Encyclopedia of Genes and Genomes (KEGG) [75].

Significant functions were represented in the form of Upset plots [76] from which the number of functional elements specific and shared by each breast cancer subtypes can be observed —see Figure 5E—.

### 4.5. Methylated Genes Meta-Analysis

Individual genes were identified from the mapping of the DMRs. Those genes with a common differential pattern between case and control groups were selected. For each gene, the p-values of all comparisons were then integrated using the Fisher combination method (or the inverse normal/weighted method). Finally, the combined p-values were corrected by the FDR method [73]. This meta-analysis strategy provided the genes with a significant common methylation profile in all the selected studies.

### 4.6. Functional DNA Methylation Signatures Meta-Analysis

Finally, once the functional enrichment study was performed for each comparison, the results were integrated into a functional meta-analysis as previously described [17]. The metafor R package was used to assess the combined effect of the studies together with a random effects model [77]. The variability of individual studies was considered for the calculation of the log odds ratio (LOR) in the meta-analysis. In turn, an analysis of heterogeneity to check the suitability of the selected studies, together with a sensitivity analysis and assessment of bias to detect whether any of the comparisons had an excessive influence on the final meta-analysis were performed.

Each analyzed function in the meta-analysis is accompanied by the combined estimate of the effect of the studies (LOR), the 95% confidence interval and the adjusted p-value by the Benjamini and Hochberg method [73]. Thus, those functions with an adjusted p-value equal to or less than 0.05 were considered significant. For each significant function, forest and funnel plots were used to measure the contribution of each study to the meta-analysis and to assess its variability —Figures 5F, 5G—.

### 4.7. Web Tool

The large volume of data and results generated in this study is freely available in the metafun-BC web tool (https://bioinfo.cipf.es/metafun-BC), which will allow users to review the results described in the manuscript and any other results of interest to researchers. The front-end was developed using the Bootstrap library. All graphics used in this tool were implemented with Plot.ly, except for the exploratory analysis cluster plot, which was generated with the ggplot2 R package.

This easy-to-use resource is organized into five sections: (1) a quick summary of the results obtained with the analysis pipeline in each of the phases. Then, for each of the studies, the detailed results of (2) the exploratory analysis, (3) the differential expression, and (4) the functional characterization are shown. The user can interact with the tool through its graphics and search for specific information for a gene or function. Finally, in Section (5), indicators are shown for the significant functions and genes identified in the methylation meta-analysis that inform whether they are more or less active in patients. Clicking on each indicator obtains the forest plot and funnel plot that explain the effect of each function in individual studies, as well as an evaluation of their variability.

## 5. Conclusions

Breast cancer is one of the main focuses of research at present, being key to the identification of biomarkers that impact on the knowledge of the disease as well as on the development of diagnostic and predictive tools that would improve the clinical decision making. In this study we have identified a series of new methylation patterns common in the different breast cancer subtypes through the application of a novel methodology based on meta-analysis tools at gene and functional level, integrating the information described so far. This approach provides greater statistical power than individual studies, incorporating in the statistical model the specific characteristics of each study. Among the main functions shared by the different breast cancer subtypes are the overactivation of the immune system itself in favor of the creation of a tumor microenvironment. Although further studies are required to fully verify and explore these findings, our results provide new clues to understand the molecular mechanisms in breast cancer.

## Author Contributions

Conceptualization, F.G.-G.; Data curation, A.M.T.-F., A.V.-M., P.M.-M. and S.R.-G.; Formal analysis, A.M.T.-F., P.M.-M. and S.R.-G.; Funding acquisition, F.G.-G; Investigation, A.M.T.-F., Z.A., D.C. and F.G.-G.; Methodology, D.C. and F.G.-G.; Project administration, F.G.-G.; Software, A.M.T.-F., P.M.-M. and S.R.-G.; Supervision, F.G.-G.; Validation, A.M.T.-F.; Visualization, A.M.T.-F., P.M.-M., S.R.-G. and F.G.-G.; Writing – original draft, M.R.H., B.G.-C., M.P.G., A.M.T.-F., Z.A., R.S.-B., A.R., J.A.L.-G., M.d.l.I.-V. and F.G.-G.; Writing – review & editing, A.V.-M., B.D.R., N.R.D., A.M.T.-F., Z.A., D.C., R.S.-B., A.R., J.A.L.-G., H.G.-M.; M.d.l.I.-V. and F.G.-G. All authors have read and agreed to the published version of the manuscript.

## Acknowledgments

The authors thank the Principe Felipe Research Center (CIPF) for providing access to the cluster, co-funded by European Regional Development Funds (FEDER) in Valencian Community 2014-2020. The authors also thank Stuart P. Atkinson for reviewing the manuscript.

## Abbreviations

The following abbreviations are used in this manuscript:

Lum-A: Luminal A
Lum-B: Luminal B
TLA: Three letter acronym
LD: linear dichroism
TNBC: Triple negative breast cancer
IDC: Invasive ductal carcinoma
ILC: Invasive lobular carcinoma

## Appendix A. Supplementary material

### Appendix A.1. Data availability

Data used in this work can be downloaded at GEO: GSE52865, GSE59901, GSE101443, GSE141338.

### Appendix A.2. Computed code

The code used in this work can be found at https://github.com/atrassierra/methylation_meta-analysis

